# Target-enrichment sequencing yields valuable genomic data for difficult-to-culture bacteria of public health importance

**DOI:** 10.1101/2022.02.16.480634

**Authors:** Tristan P. W. Dennis, Barbara K. Mable, Brian Brunelle, Alison Devault, Ryan Carter, Clare L. Ling, Blandina T. Mmbaga, Jo E. B. Halliday, Katarina Oravcova, Taya L. Forde

## Abstract

Genomic data contribute invaluable information to the epidemiological investigation of pathogens of public health importance. However, whole genome sequencing (WGS) of bacteria typically relies on culture, which represents a major hurdle for generating such data for a wide range of species for which culture is challenging. In this study, we assessed the use of culture-free target-enrichment sequencing as a method for generating genomic data for two bacterial species: 1) *Bacillus anthracis,* which causes anthrax in both people and animals and whose culture requires high level containment facilities; and 2) *Mycoplasma amphoriforme*, a fastidious emerging human respiratory pathogen. We obtained high quality genomic data for both species directly from clinical samples, with sufficient coverage (>15X) for confident variant calling over at least 80% of the baited genomes for over two thirds of the samples tested. Higher qPCR cycle threshold (*Ct*) values (indicative of lower pathogen concentrations in the samples), pooling libraries prior to capture, and lower captured library concentration were all statistically associated with lower capture efficiency. The *Ct* value had the highest predictive value, explaining 52% of the variation in capture efficiency. Samples with *Ct* values ≤ 30 were over 6 times more likely to achieve the threshold coverage than those with a *Ct* > 30. We conclude that target-enrichment sequencing provides a valuable alternative to standard WGS following bacterial culture and creates opportunities for an improved understanding of the epidemiology and evolution of many clinically important pathogens for which culture is challenging.

**Data summary:** The authors confirm all supporting data, code and protocols have been provided within the article or through supplementary data files. Scripts used in this study can be accessed on GitHub at https://github.com/tristanpwdennis/bactocap. All sequence data generated during this study have been deposited in the European Nucleotide Archive (ENA) Sequence Read Archive (SRA) under project accession numbers PRJEB46822 (*B. anthracis*) and PRJEB50216 (*M. amphoriforme*). Accession numbers for individual samples, along with metadata, laboratory parameters and sequence quality metrics, are available at the University of Glasgow’s data repository, Enlighten, at http://dx.doi.org/10.5525/gla.researchdata.1249.

## Introduction

Genomic data can provide critical information towards understanding bacterial epidemiology and evolution, and thereby help to guide public health policy and infectious disease control [1]. Whole genome sequence (WGS) data contribute to determining transmission dynamics at various scales [2], and are used to track the emergence and evolution of particular strains or traits [3]. WGS is routinely implemented for the characterisation of a wide range of bacteria, but typically involves working from cultured isolates to yield DNA of sufficient quantity and quality. This reliance on culture as a first step represents a major hurdle for generating genomic data for many bacteria of public health importance. For instance, for bacteria with high biosafety requirements, suitable laboratories may not be available. Similar obstacles exist for fastidious bacteria (i.e. due to their complex nutritional requirements and slow growth), rendering their culture for genetic analysis difficult or impossible [4]. Moreover, while sequencing can be performed directly on extracts from clinical samples (e.g. bodily fluids and tissues), pathogen DNA is often present only in low quantity, representing only a fraction of the overall DNA in a sample [5], thus complicating WGS-based analyses.

Hybridization capture followed by sequencing, or ‘target-enrichment sequencing’, is one approach that may be able to overcome this obstacle and enable bacterial genomic sequence data to be obtained more effectively from clinical samples [6, 7]. In this culture-free method, libraries are prepared from DNA extracted from the sample, after which specific fragments are selectively enriched prior to sequencing. This is accomplished using custom-designed probes, or ‘baits’ - typically RNA oligonucleotides labelled with biotin that hybridize to the DNA of interest and are then captured by streptavidin-coated magnetic beads, allowing non-target DNA to be removed through wash steps [8]. Baits are able to selectively capture a wide range of fragment sizes [8, 9] and also allow for polymorphisms between their sequence and that of the enriched fragment; depending on the stringency of capture, this can be as low as 70% sequence similarity [8]. Target-enrichment sequencing increases the potential to reconstruct whole bacterial genome sequences from relatively small amounts of target DNA from clinical samples [4, 10–12], thereby facilitating in-depth investigations, including phylogenomics, into these otherwise difficult-to-study pathogens.

In this study, we evaluated target-enrichment sequencing as a means of generating genomic data directly from samples containing two different difficult-to-culture bacteria: *Bacillus anthracis* and *Mycoplasma amphoriforme. Bacillus anthracis* is a Gram positive bacterium that causes anthrax – a neglected bacterial zoonosis that is a threat to livestock and human health in low-and-middle-income countries (LMICs) [13]. The high virulence of *B. anthracis* makes it hazardous to culture and therefore requires containment level (CL) 2+ or 3 facilities [14]. Limited high-containment laboratory capacity in LMICs thus represents a barrier to the generation of sequence data for *B. anthracis* and understanding of its diversity and transmission in endemic settings. Moreover, concerns over the use of *B. anthracis* as a biological weapon make the shipment of samples for culture elsewhere challenging [15]. Although the true clinical significance of *M. amphoriforme* is yet to be elucidated, it has been associated with respiratory tract infections in immunocompetent and immunocompromised patients [16–21]. While there is great potential to use comparative genomics to gain insights into its pathogenicity and epidemiology, *M. amphoriforme* requires specialised culture media and conditions; several weeks are required to generate bacterial cultures with suitable cell densities for DNA extraction and not all PCR positive samples yield a positive culture despite high estimated bacterial loads [16, 18].

The primary objective of this study was to assess the feasibility of generating sequence data for *B. anthracis* and *M. amphoriforme* at coverage suitable for variant calling over the majority of the genome. We also sought to determine the upper pathogen-specific qPCR cycle threshold (*Ct*) value limit to provide sufficient coverage for accurate variant calling, and the influence of different variables (i.e. *Ct*, whether libraries were pooled prior to bait capture, and concentration of the captured libraries) on capture efficiency (i.e. the proportion of on-target reads based on reference mapping).

## Methods

Custom myBaits^®^ sets were designed for *B. anthracis* and *M. amphoriforme* in partnership with Daicel Arbor Biosciences (Ann Arbor, Michigan, USA). RNA baits of ~80 base pairs (bp) were designed to achieve 50% overlap (tiling) across targeted regions (i.e. at least two baits covering any given position). For *B. anthracis,* baits were designed to tile the core chromosomal genome. Publicly available *B. anthracis* genomes available at the time of bait design (n=52; Table S1) – including genomes from all three major genetic subgroups (Clades A, B and C) [22] – were used to generate a core genome alignment in Parsnp [23]. This alignment was 4.66 MB (89% of the 5.23 MB Ames Ancestor reference chromosomal genome, NC_007530.2 [24]). For *M. amphoriforme*, bait sequences were designed to cover the pangenome, which was constructed using all *M. amphoriforme* whole genome sequences available at the time of the experimental design, and included the A39 reference genome NCTC 11740 (accession number HG937516; 1,029,022 bp) and 18 additional *M. amphoriforme* genomes [19]. Pairwise comparison of these genomes was then done using Artemis Comparison Tool [25], concatenating any regions of difference to the reference genome of strain A39, resulting in a pangenome of 1,064,467 bp. For both bacterial species, baits that targeted loci with high copy number in the genome were omitted (e.g. rRNA, tRNA), as were simple repeats, which were soft-masked using RepeatMasker [26]. Additionally, *in silico* analysis was performed to identify and remove any baits that could potentially cross-hybridize with host or non-target bacteria (File S1, Table S2).

For *B. anthracis*, DNA extracts (n = 93) from animal carcasses sampled in northern Tanzania between 2016 and 2018 were analysed. All sample extracts had tested positive by qPCR for three regions of the *B. anthracis* genome (*cap, lef* and *PL3* [27]), with one exception where *cap* failed to amplify. Taking the highest of the three *Ct* values per sample, the *Ct* values ranged from 19 to 37 (median = 25). For *M. amphoriforme*, DNA extractions from nasopharyngeal swabs (NPS, n=52) and nasopharyngeal aspirates (NPA, n=4) that had tested positive with a *M. amphoriforme-*specific qPCR targeting the *udg* gene were included [18]. The NPS originated from a longitudinal *Streptococcus pneumoniae* carriage study on the Thailand-Myanmar border [28], and were collected from 11 participants, each providing between three and six consecutive swabs between 2007-2010. The NPA originated from influenza virus PCR surveillance also on the Thailand-Myanmar border in 2016 and 2017. The *Ct* values for *M. amphoriforme* samples ranged from 20 to 40 (median = 29). Sample-specific metadata for both bacterial species are available as Supplementary Tables A & B at the University of Glasgow’s Enlighten data repository [29].

All DNA extracts were quantified using Qubit dsDNA high sensitivity Assay Kits (Thermo Fisher Scientific). All initial libraries (referred to as “pre-capture libraries”) were prepared using the NEBNext^®^ Ultra II FS DNA Library Prep Kit for Illumina with NEBNext Sample Purification Beads (New England BioLabs). This protocol includes a short PCR amplification (5-7 cycles) to add sample-specific barcodes and to increase the total DNA concentration prior to bait capture. We varied the number of cycles across library preparation batches to achieve the recommended minimum DNA input for bait capture (100 ng), whilst aiming to use the minimum number of cycles required (details in File S1).

For bait capture, we followed the myBaits^®^ Hybridization Capture for Targeted NGS protocol (Daicel Arbor Biosciences, Manual v4.01). For each bait capture, 7 μl of pre-capture library was used as input. Most libraries were captured individually. For a subset, libraries were pooled before bait hybridisation with the goal of reducing the cost of baits per sample; this was done for 34 *B. anthracis* libraries that spanned a range of *Ct* values, and eight *M. amphoriforme* libraries prepared from samples with higher *Ct* values. As per the myBaits^®^ protocol, we aimed to include four libraries per pool; however, in some cases where certain initial library preparations were unsuccessful, only three pre-capture libraries were pooled. For each pool, approximately equal input concentrations of each pre-capture library were included to yield a total input of 125 ng in 7 μl.

Following bait hybridisation, the number of cycles in the PCR amplification step – included in the myBaits^®^ protocol to yield at least 1.5 ng/μl DNA for downstream sequencing – was varied among preparation batches; between 12 and 15 cycles were used for *B. anthracis* and 12 or 13 cycles for *M. amphoriforme* samples. Post-amplification clean-up was done by column-based purification using Zymo DNA Clean & Concentrator −5 kit (Cambridge Bioscience, UK), eluting in 12 μl of 0.1X TE buffer. We refer to the final product as ‘captured libraries’.

Captured libraries were quality checked using a Bioanalyzer (Agilent) and sequenced at Glasgow Polyomics on an Illumina NextSeq instrument, generating 75 bp paired-end reads. Given differences in genome size, approximately 4 and 1 million reads were targeted for *B. anthracis* and *M. amphoriforme,* respectively. Raw reads were trimmed with fastp 0.20.1 with default settings [30] and aligned to the *B. anthracis* Ames Ancestor reference genome or the *M. amphoriforme* pangenome, respectively, with bwa-mem v0.7.17-r1188 [31], after which PCR duplicates were removed in GATK [32]. We defined a ‘successful’ sequencing outcome as a sample achieving >15X coverage (the cut-off for confidently calling a genotype in a haploid organism [33]) over 80% of the baited genome (i.e. the parts of the core *B. anthracis* genome or *M. amphoriforme* pangenome against which baits were designed). Full analysis scripts are available on Github [34].

A binomial generalised linear mixed model (glmm) was performed in R [35] with *glmer* in the *lme4* library [36] to investigate the association between the capture efficiency – defined as the proportion of mapped reads per sample – and the explanatory variables: *Ct* values from pathogen-specific qPCR performed on original sample DNA extracts; bacterial species (i.e. *B. anthracis* or *M. amphoriforme*); whether or not libraries were pooled; and captured library concentration (ng/μl). All models included an observation level random effect to account for overdispersion [37]. Pairwise interactions between bacterial species and the other variables were also considered. Model simplification was evaluated based on comparison of AIC, pseudo-R2 analysis was performed in the *MuMIn* library [38], and the final model was plotted in *sjPlot* [39].

## Results

The final bait set for *B. anthracis* comprised 148,811 baits and covered 4,637,856 bp (88%) of the 5,227,419 Ames Ancestor chromosomal reference sequence. For *M. amphoriforme*, the final bait set comprised 24,444 baits. This set covered 1,002,426 bp (97%) of the 1,029,022 bp A39 reference genome, or 94% of the 1,064,467 bp pangenome. Final bait set sequences for both case studies (as .bed annotations) are provided on GitHub [34], and are also available through the Daicel Arbor Biosciences Community Panels webpage.

For *B. anthracis*, the median number of total reads generated per sample was 6,413,796 (range: 539,623 - 32,708,157). The mean depth of coverage per sample across the baited regions of the Ames Ancestor reference genome ranged from 0.2X to 449.4X (median = 43.2X) (Fig. 1A). Based on our previously defined threshold of >15X coverage over >80% of the baited genome, 55 of 93 *B. anthracis* samples (59.1%) were considered successfully sequenced: 40/59 (68%) of unpooled and 15/34 (44%) pooled samples. The proportion of the baited genome covered to 15X depth ranged from 0-100% per sample (median = 97.3%; Fig. 1B). The proportion of mapped reads per sample ranged from 2.3% to 99.2% (median = 69.7%) (Fig. 1C). This latter metric is independent of the per-sample sequencing depth.

**Fig. 1.**
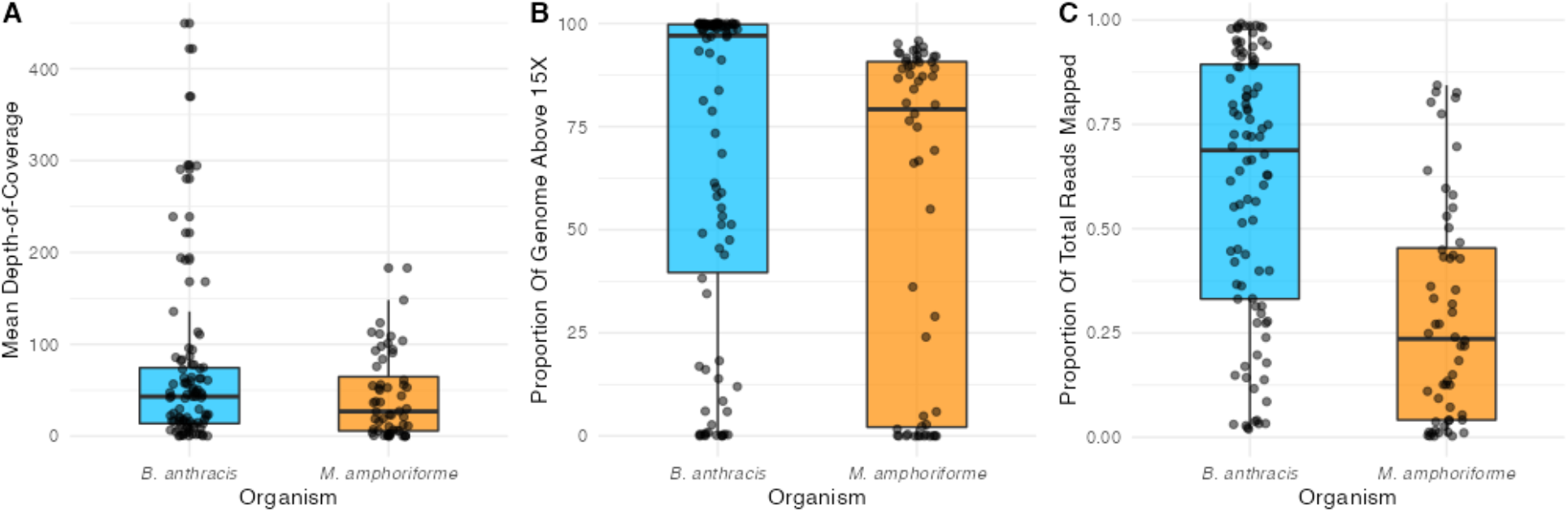
Target-enrichment sequencing outcomes. A) Mean depth-of-coverage over baited regions of the reference genomes, per-sample for *B. anthracis* (blue) and *M. amphoriforme* (orange); B) proportion of baited genome covered to > 15X per-sample; and C) proportion of reads mapped to baited regions. Points indicate individual samples, box and whisker plots indicate maximum, minimum, upper-lower quartiles and median values.

For the *M. amphoriforme*-positive samples, the total number of reads generated per sample ranged from 1,082,454 to 4,247,924 (median = 2,989,985). The mean depth of coverage per sample across the baited regions of the *M. amphoriforme* pangenome ranged from 0.1X to 183.1X (median = 27.0X) (Fig. 1A). Twenty-eight of 56 samples (50%) achieved >15X coverage over 80% of the baited *M. amphoriforme* pangenome: 28/48 (58%) of unpooled and 0/8 (0%) pooled. The proportion of the pangenome covered to this depth ranged from 0 - 95.8% (median = 79.2%; Fig. 1B). Mapped reads represented between 0.3% and 84.4% of the total reads per sample (median = 23.6%) (Fig. 1C). Summary statistics of the sequencing metrics for both datasets are provided in Table S3. Sample-specific sequencing results and accession numbers for both bacterial species are available in the Supplementary Tables on Enlighten [29].

The final model to evaluate variables influencing capture efficiency included the *Ct* value, pooling libraries before capture (yes/no), and captured library concentration variables, with an observation level random effect (Table S4; File S1). The model indicated that lower *Ct* is significantly associated with increased capture efficiency (Fig. 2). Captured library concentration was significantly positively associated with capture efficiency (Fig. S1), while pooling libraries prior to bait capture was negatively associated with capture efficiency (Fig. S2). The model accounted for 55% of the variation in capture efficiency. We found that *Ct* explained 52% of the variation in our data. In contrast, captured library concentration explained 13%, and pooling only 1%. Of 10 *B. anthracis* samples with *Ct* > 30, only one of these was successfully sequenced (with a *Ct* of 31) vs. 65% of samples (54/83) with *Ct* ≤ 30. Similarly for *M. amphoriforme,* 25 of 33 samples with *Ct* ≤ 30 (76%) were successful, compared with 3 of 23 with *Ct* > 30 (13%).

**Fig. 2.**
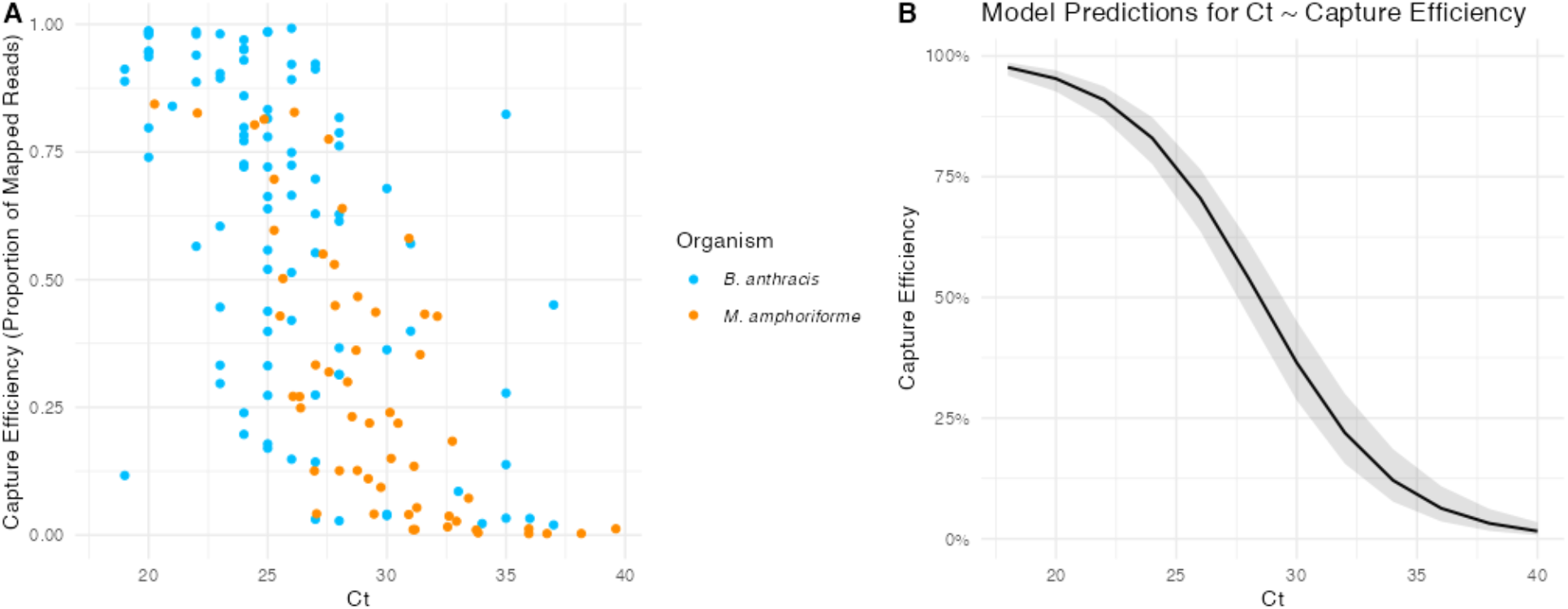
Relationship between *Ct* value and capture efficiency. A) Each point represents an individual sample, with bacterial species distinguished by colour. B) Binomial glm model prediction of the relationship between *Ct* value and capture efficiency for both species. Shading represents a 95% prediction interval.

## Discussion

We have shown that broad, deep sequencing coverage can be obtained for *M. amphoriforme* and *B. anthracis* directly from samples using target-enrichment sequencing. These species are representative of the breadth of typical bacterial genome sizes (~1.0 MB and ~5.2 MB, respectively). For both species, we were able to recover a substantial portion of the genome at a sufficient depth of coverage for haploid variant calling where otherwise, in the experience of the authors, such data would have been extremely challenging or impossible to obtain. Since total sequencing read amounts are independent of enrichment performance, it is possible that even more of the study samples could have reached our coverage threshold (>15X across 80% of the baited genome) had we performed additional sequencing to generate more reads per sample. Target-enrichment sequencing will be transformative for the molecular epidemiological study of these two species of public health importance, and more generally provides a valuable alternative when standard WGS following bacterial culture is impracticable [11, 12, 40, 41], so long as a reference sequence is available to facilitate bait design. Culture-free methods such as targeted-sequence capture expand the opportunities for the genomic study of several Hazard Group 2+/3 pathogens, such as *B. anthracis*, within LMIC countries where they remain a threat to human and animal health. Whereas high-containment facilities are often lacking, sequencing platforms are increasingly available worldwide. Targeted-sequence capture also promises to facilitate our understanding of the genomic epidemiology of *M. amphoriforme,* which has only been isolated from patients on a few occasions due to its requirement for specific culture conditions and slow growth [19]. It has particular promise for studying pathogenic and antimicrobial resistance determinants, and thus aid timely clinical management of respiratory tract infections caused by this bacterium [17, 19, 21]. The developed bait sets and methods described in this study will provide a valuable resource for other researchers wishing to pursue the genomic study of these bacteria.

Our results suggest that target-enrichment sequencing is most effective for samples with higher pathogen-specific DNA concentrations, as indicated by *Ct* value. *Ct* was strongly correlated with capture efficiency (i.e. a higher proportion of mapped reads, indicative of less off-target capture), as well as the success of sequencing to a sufficient depth for confident genotyping, a finding consistent with previous studies [11, 42]. At least two thirds of samples with *Ct* values ≤ 30 – indicative of higher starting pathogen concentrations in the samples – were successfully sequenced for both sample sets, whereas this success rate dropped to 10-13% for samples with *Ct* > 30. Based on previous studies implementing the *udg* gene as a PCR target for *M. amphoriforme* [18], a *Ct* of 30 would correspond to approximately 90-900 genome copies per PCR reaction, suggesting that target-enrichment sequencing will be most useful for pathogens that are present in moderate quantities in clinical samples. Generating genomic data for clinical samples with very low pathogen concentrations thus remains a challenge. While not assessed in this study, including a second round of bait capture (i.e. putting the enriched library through a second round of enrichment) has been suggested to improve sensitivity for such samples [43]. Although capture efficiency was also significantly associated with higher captured library concentration and with libraries not having been pooled prior to capture, the predictive value of these variables was low; *Ct* value alone could explain most of the variation in the data.

We found that pooling up to four libraries for capture still yielded the targeted coverage in nearly half of the pooled *B. anthracis* samples, thereby representing potential for cost savings by reducing the reagents associated with bait capture. The *M. amphoriforme* samples with libraries pooled in this study had a higher median *Ct* than unpooled samples (File S1), which likely accounts for the greater difference in capture efficiency observed between pooled vs unpooled libraries for the two bacterial species (Fig S2). Previous studies have been more consistent in achieving successful coverage when pooling as many as 20 libraries for *Borellia burgdorferi* [10] and *Treponema pallidum* [40]; this success may be attributed in part to carefully selecting libraries for pooling within a narrow range of similar *Ct* values as performed by Beale *et al*. [40] and as recommended in later versions of the MyBaits^®^ manual (v5).

The ability to generate culture-free genomic data opens significant opportunities for wider phylogenetic studies of these two bacterial species of public health concern, and highlights the potential value of target-enrichment sequencing for facilitating an improved understanding of the epidemiology and evolution of many clinically important pathogens for which genomic data have been limited.

## Author Statements

### Author contributions

Conceptualisation: BKM, JEBH, KO, TLF. Data curation: TPWD, KO, TLF. Formal analysis: TPWD, KO, TLF. Funding acquisition: JEBH, BKM, KO, TLF, BB, AD. Investigation: RC, KO, TLF. Methodology: BB, AD, BKM, JEBH, TPWD, KO, TLF. Project administration: JEBH, BKM, KO, TLF. Resources: BTM, CLL. Software: TPWD. Visualisation: TPWD. Writing – original draft: TPWD, KO, TLF. Writing – review & editing: TPWD, BKM, BB, AD, CLL, JEBH, KO, TLF.

### Research and ethical approval

The *B. anthracis* study received ethical approval from the Kilimanjaro Christian Medical University College Ethics Review Committee (certificate No. 2050); the National Institute for Medical Research, Tanzania (NIMR/HQ/R.8a/Vol. IX/2660); Tanzanian Commission for Science and Technology (2016-95-NA-2016-45); and the College of Medical Veterinary and Life Sciences ethics committee at the University of Glasgow (200150152). Material and data transfer agreements were established as part of these approvals, and this study complied with Tanzania’s national access measures for genetic material. Ethical approval related to the *M. amphoriforme* samples was granted in the original study by ethics committees of the Faculty of Tropical Medicine, Mahidol University, Thailand (MUTM-2009-306) and Oxford University, UK (OXTREC-031-06) [23].

## Acknowledgements

We are grateful to the members of the field team who collected the *B. anthracis* samples: Deogratius Mshanga, Sabore Ole Moko, Sironga Nanjicho and Kadogo Lerimba. We also thank Nichith Kollanandi Ratheesh and the staff at Kilimanjaro Clinical Research Institute for assistance in sample processing, particularly Alutu Masokoto. Thank you to Shoklo Malaria Research Unit, Thailand for the provision of *M. amphoriforme* positive samples, in particular Prof Paul Turner and Prof François Nosten. We would like to thank Dr Mat Beale and Prof Nick Thomson from Wellcome Sanger Institute for their invaluable input for *M. amphoriforme* data analysis and constructive suggestions for the manuscript. We also thank Dr Paul Johnson and Prof Dan Haydon for their input on the statistical analyses.

## Supplementary Information

**File S1. Supplementary Methods and Results.** Full details of the bait set design, sample processing, library preparation, and additional bioinformatics steps, including the generation of mapping and coverage statistics.

**Fig S1.** Relationship between captured library concentration and capture efficiency for both species. Shading represents a 95% prediction interval.

**Fig. S2.** Box-and-whisker plots overlaid with jittered points indicating capture efficiency vs whether the sample was pooled or unpooled prior to bait capture. Shown for both *B. anthracis* and *M. amphoriforme*.

**Table S1.** Publicly available *Bacillus anthracis* genome sequences used to generate a core genome alignment for the design of *B. anthracis* specific baits.

**Table S2.** Human respiratory commensal and pathogenic bacterial genome sequences used for the design of *M. amphoriforme* specific baits. Non-specific baits with BLAST hits to these organisms were removed.

**Table S3. Summary statistics.** Total reads, mapped reads, fraction of mapped reads, mean depth of coverage (doc), fraction of the genome covered > 15X, for *B. anthracis* and *M. amphoriforme.*

**Table S4.** Model details for binomial glmm of the effect of *Ct* value, captured library concentration and pooling prior to bait capture on capture efficiency, including an individual-level random effect.

