## Supplementary material for "Target-enrichment sequencing yields valuable genomic data for difficult-to-culture bacteria of public health importance"

### File S1

#### Supplementary Methods

##### *Bait Design*

For *B. anthracis*, approximately 6 million baits were initially designed; however, after removal of 100% duplicate bait sequences, 2.2 million baits remained, which were queried against the Ames ancestor reference genome using BLASTn to verify hits and remove those that might have been designed across artificial 'joints' in the core genome. This resulted in the removal of 12,796 baits with the longest BLAST hit < 65 bp. To minimize over-tiling towards the 3' end of the alignment (due to staggered bait start positions), baits with 100% identity across more than 75% overlap were clustered, with only one bait per cluster retained (n = 161,899 baits). BLAST queries against targeted mammalian genomes (human, cow, sheep, goat, pig, mouse, rat, camel and donkey) led to the removal of only 3 baits with hits. Baits with >50% simple repeat masking (n=314) were also removed from the bait set. Baits overlapping with any tRNA or rRNA annotations in the reference sequence (n = 171) were removed, as these would be expected to be less specific for *B. anthracis* (i.e. conserved across various bacterial species). Remaining baits were queried against a local database of NCBI RefSeq bacterial genomes using BLASTn, and results parsed with MEGAN6 [1] (min bit score 70, top % 100, min support 0%); baits were removed for any taxonomic rank higher than class Bacilli (i.e. less specific), thus removing a further 1266 baits. Finally, baits with > 98% identity across > 75% overlap were clustered, and 1 bait retained per cluster, effectively removing baits with 1 SNP difference. For *M. amphoriforme*, a total of 25,961 raw baits were designed. Each bait candidate was aligned *in silico* against relevant commensal and pathogenic bacterial genomes (potentially residing in the human respiratory tract and oropharynx; Table S2) and the human genome using BLAST. Baits with hits to these bacterial and human genomes were removed to avoid potential cross-hybridization with exogenous DNA from non-target organisms. Baits were also filtered that were > 25% soft masked for simple repeats.

##### *Sample selection and processing*

For *B. anthracis*, samples were stored for up to several months at ambient temperature prior to DNA extraction. DNA was extracted from various sample types using the DNeasy Blood & Tissue kit (Qiagen, UK), with minor modifications to initial sample preparation depending on the sample type as previously described [2]. Final extracts were filtered using a 0.2 µm spin column (Corning Life Sciences) to remove any potentially viable spores prior to downstream molecular analyses. Extracts in this study were from tissues (n = 52), blood swabs (n = 30), whole blood (n

= 9) or insects collected on the carcass (n = 2). Samples were primarily from livestock species, namely sheep (n = 61), cattle (n = 9), goats (n = 5) and a donkey (n = 1), although some samples from wildlife were also included, namely wildebeest (n = 8) and zebras (n = 7); species was not recorded for two samples. All nasopharyngeal swab (NPS) samples collected for *M. amphoriforme* testing were stored at -80 °C. DNA was extracted using Wizard DNA extraction kit (Promega) according to manufacturer's instructions. Sample-specific metadata for both bacterial species are available as Supplementary Tables A & B at University of Glasgow's Enlighten data repository [3].

***Library preparation and bait capture***

Samples were prepared in batches of up to 28 samples. Variations made to any experimental parameters are detailed in the supplementary tables on Enlighten [3]. For *B. anthracis*, libraries were prepared following either the Chapter 1 protocol (for inputs ≤ 100 ng) or the Chapter 2 protocol (for inputs ≥ 100 ng) of the NEBNext® Ultra II FS DNA Library Prep Kit; Chapter 1 was followed for all *M. amphoriforme* samples. If initial concentrations of DNA extracts were higher than recommended for the library preparation protocols (i.e. > 100 ng if using Chapter 1, or > 500 ng if using Chapter 2), DNA was diluted in 0.1X TE buffer to achieve the recommended maximum. Dual index NEBNext Multiplex Oligos were used throughout, with the exception that single index primers were used during a trial run with one *B. anthracis* sample. Enzymatic fragmentation was done by incubating at 37°C for 8 minutes, except for *B. anthracis* library batch 5, where incubation was for 10 minutes. For *B. anthracis* NEB library batch 1, 5 amplification cycles were used. Since this resulted in a high proportion of pre-capture libraries with concentrations below the recommended input for bait capture, i.e. 100 ng in 7 µl, from Batch 2 onwards 6 amplification cycles were used, except for Batch 5, where this was increased to 7 cycles. For *M. amphoriforme,* all libraries were prepared using 6 amplification cycles. For *B. anthracis* library batch 1, pre-capture library DNA was eluted in 30 µl of 0.1X TE buffer; from batch 2 onwards, elution was in 23 µl to increase the concentration. For some pre-capture libraries with low initial concentrations as measured on Qubit, libraries were further concentrated prior to bait capture using the Zymo DNA Clean & Concentrator -5 kit (Cambridge Bioscience, UK) enabling elution in very small volumes (i.e. 10 µl). Some captured libraries in *B. anthracis* batches 1 and 2 (in which only 12 amplification cycles were implemented post-hybridisation) had very low concentration, and were therefore subjected to a further 5 PCR amplification cycles, followed by a second clean-up step.

***Bioinformatics***

After trimmed reads were mapped using bwa-mem, duplicate reads were removed, and read groups added with *gatk* v4.2.1.0 MarkDuplicates and AddOrReplaceReadGroups as part of their standard pipeline [4]. Mapping and coverage statistics were collected by calculating coverage over baited regions using bait .bed annotation files in *gatk DepthOfCoverage*. Mean depth-of-coverage was calculated as: number mapped reads * read length / length of the baited reference genome. Samtools v1.3.1 *flagstat* was used to collect data on the number of duplicate reads [5]. The workflow was implemented in *nextflow* v20.10.0.5430 [6]. Scripts for the analysis of both datasets were written in R [7], incorporating *tidyverse* v1.3.0 [8], *DHARMa* v0.4.3 [9] and *cowplot* v1.1.1 [10]. The *nextflow* workflow, Docker container link, and analysis scripts are available at https://github.com/tristanpwdennis/bactocap.

### Supplementary Results

#### *Ct values of sequenced samples*

Taking the highest of the *Ct* values per sample from qPCR of the three *B. anthracis* genomic targets, the median was 25 across the 93 samples; this was the same for both the group of pooled and unpooled samples. The variance in *Ct* for unpooled samples was 21.9, and for pooled samples was 4.9. The median (and variance) of *Ct* values of qPCR from *M. amphoriforme* was 29 (8.3), whereas it was 36 (5.6) for pooled samples. Sample-specific sequencing results are available on Enlighten [3].

#### *Captured library concentrations*

Following bait capture, concentrations of captured *B. anthracis* libraries ranged from 0.16 to 66 ng/µl, with a median post-capture library concentration of 1.6 ng/µl (variance = 166.3). For *M. amphoriforme,* captured library concentrations ranged from 0.2 to 3.3 ng/µl, with a median of 0.8 ng/µl (variance = 0.58).

### Supplementary Figures

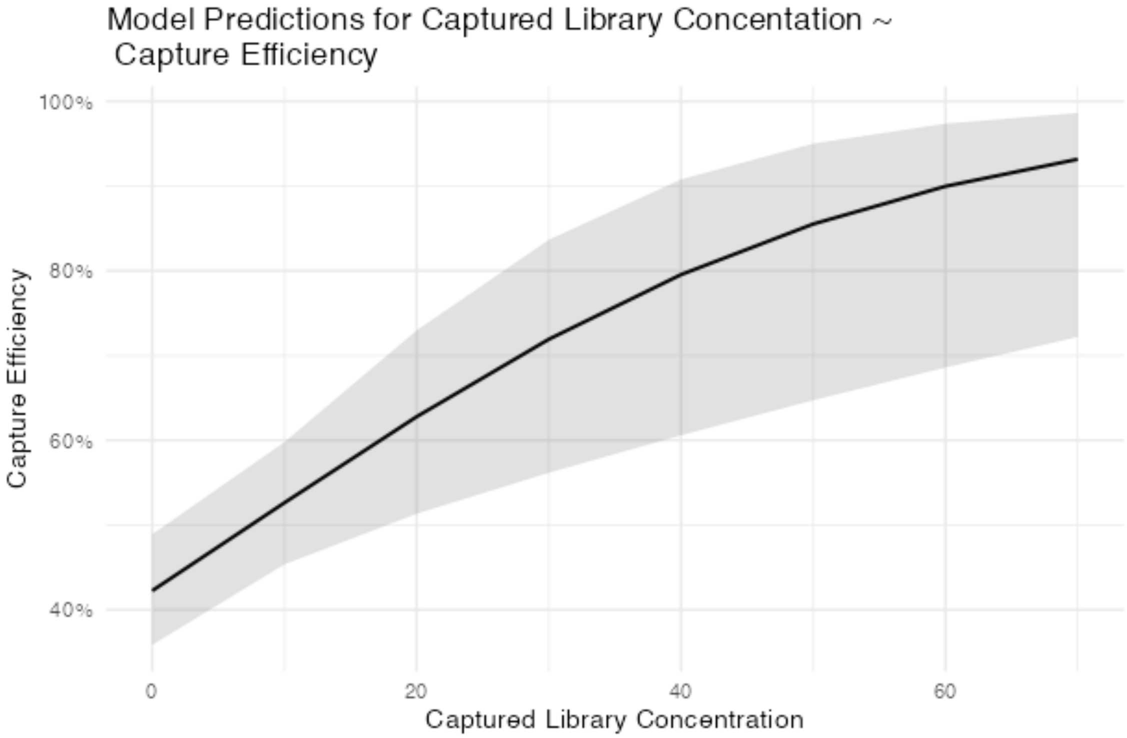


**Fig. S1.** Relationship between captured library concentration and capture efficiency for both species. Shading represents a 95% prediction interval.

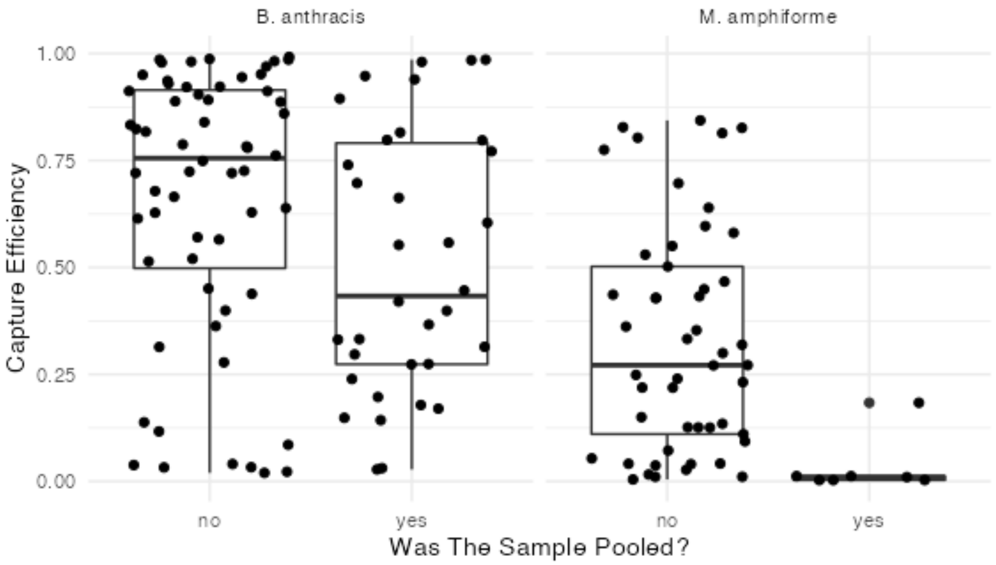


**Fig. S2.** Box-and-whisker plots overlaid with jittered points indicating capture efficiency vs whether the sample was pooled or unpooled prior to bait capture. Shown for both *B. anthracis* and *M. amphoriforme.*

### Supplementary Tables

**Table S1. Publicly available *Bacillus anthracis* genome sequences used to generate a core genome alignment for the design of *B. anthracis* specific baits.**

| Strain | BioSample | BioProject | Assembly | NCBI Reference Sequence |
|---|---|---|---|---|
| 2000031021 | SAMN02736984 | PRJNA243523 | GCA_000742655.1 | NZ_CP007618.1/CP007618.1 |
| 2002013094 | SAMN03174509 | PRJNA238050 | GCA_000832965.1 | NZ_CP009902.1/CP009902.1 |
| HYU01 | SAMN02874036 | PRJNA231762 | GCA_000725325.1 | NZ_CP008846.1/CP008846.1 |
| SVA11 | SAMN03081486 | PRJNA217316 | GCA_000583105.1 | NZ_CP006742.1/CP006742.1 |
| BA1035 | SAMN03010427 | PRJNA238135 | GCA_000832725.1 | NZ_CP009700.1/CP009700.1 |
| RA3 | SAMN03075602 | PRJNA238136 | GCA_000832745.1 | NZ_CP009697.1/CP009697.1 |
| Tyrol 4675 | SAMN06186720 | PRJNA309927 | GCA_001936375.1 | NZ_CP018903.1/CP018903.1 |
| K3 | SAMN03010428 | PRJNA238080 | GCA_000832465.1 | NZ_CP009331.1/CP009331.1 |
| H9401 | SAMN02603474 | PRJNA49361 | GCA_000258885.1 | NC_017729.1/CP002091.1 |
| SK-102 | SAMN03012770 | PRJNA238068 | GCA_000832565.1 | NZ_CP009464.1/CP009464.1 |
| CDC 684 | SAMN02603931 | PRJNA31329 | GCA_000021445.1 | NC_012581.1/CP001215.1 |
| Vollum | SAMN02736982 | PRJNA243521 | GCA_000742895.1 | NZ_CP007666.1/CP007666.1 |
| Vollum 1B | SAMN03010433 | PRJNA238082 | GCA_000832445.1 | NZ_CP009328.1/CP009328.1 |
| Pasteur | SAMN03024436 | PRJNA238046 | GCA_000832585.1 | NZ_CP009476.1/CP009476.1 |
| Smith 1013 | SAMN02732407 | PRJNA243516 | GCA_000742315.1 | NZ_CM002879.1/CM002879.1 |
| A0157 | SAMN03267488 | PRJNA270580 | GCA_000808075.1 | NZ_CP010342.1/CP010342.1 |
| Turkey32 | SAMN03010432 | PRJNA236040 | GCA_000833275.1 | NZ_CP009315.1/CP009315.1 |
| PAK-1 | SAMN03010430 | PRJNA237808 | GCA_000832425.1 | NZ_CP009325.1/CP009325.1 |
| A1144 | SAMN02999504 | PRJNA257008 | GCA_000875715.1 | NZ_CP010852.1/CP010852.1 |
| Canadian_bison | SAMN03202901 | PRJNA238044 | GCA_000833125.1 | NZ_CP010322.1/CP010322.1 |
| Pollino | SAMN03296000 | PRJNA273788 | GCA_000831505.1 | NZ_CP010813.1/CP010813.1 |
| Larissa | SAMN03765650 | PRJNA286154 | GCA_001277955.1 | NZ_CP012519.1/CP012519.1 |
| BA1015 | SAMN03010426 | PRJNA238204 | GCA_000832665.1 | NZ_CP009544.1/CP009544.1 |
| V770-NP-1R | SAMN03092715 | PRJNA235226 | GCA_000832785.1 | NZ_CP009598.1/CP009598.1 |
| Ohio ACB | SAMN03010429 | PRJNA238205 | GCA_000832505.1 | NZ_CP009341.1/CP009341.1 |
| 52-G | SAMN02951870 | PRJNA224563 | GCA_000559005.1 | NZ_CM002395.1/CM002395.1 |

| | | | | |
|---|---|---|---|---|
| 8903-G | SAMN02951868 | PRJNA224562 | GCA_000558965.1 | NZ_CM002401.1/CM002401.1 |
| 9080-G | SAMN02951869 | PRJNA224558 | GCA_000558985.1 | NZ_CM002398.1/CM002398.1 |
| A16 | SAMN02641483 | PRJNA40303 | GCA_000512835.2 | NZ_CP001970.2/CP001970.2 |
| Ames | SAMN02603432 | PRJNA309 | GCA_000007845.1 | NC_003997.3/AE016879.1 |
| Ames Ancestor | SAMN02603433 | PRJNA10784 | GCA_000008445.1 | NC_007530.2/AE017334.2 |
| A0248 | SAMN02603932 | PRJNA33543 | GCA_000022865.1 | NC_012659.1/CP001598.1 |
| Ames | SAMN03201418 | PRJNA238045 | GCA_000833065.1 | NZ_CP009981.1/CP009981.1 |
| Stendal | SAMN04442145 | PRJNA309927 | GCA_001543225.1 | NZ_CP014179.1/CP014179.1 |
| 14RA5914 | SAMN07498358 | PRJNA397960 | GCA_002277915.1 | NZ_CP023001.1/CP023001.1 |
| Tangail-1 | SAMN05003865 | PRJNA309927 | GCA_001654475.1 | NZ_CP015779.1/CP015779.1 |
| BFV | SAMN02736972 | PRJNA243518 | GCA_000742875.1 | NZ_CP007704.1/CP007704.1 |
| Sterne | SAMN03010431 | PRJNA236483 | GCA_000832635.1 | NZ_CP009541.1/CP009541.1 |
| Sterne | SAMN02598266 | PRJNA10878 | GCA_000008165.1 | NC_005945.1/AE017225.1 |
| delta Sterne | SAMN02736981 | PRJNA243519 | GCA_000742695.1 | NZ_CP008752.1/CP008752.1 |
| Parent1 | SAMN04093743 | PRJNA295544 | GCA_001683095.1 | NZ_CP012730.1/CP012730.1 |
| SPV842_15 | SAMN06270326 | PRJNA368680 | GCA_001990245.1 | NZ_CP019588.1/CP019588.1 |
| PR02 | SAMN04075677 | PRJNA295544 | GCA_001683155.1 | NZ_CP012721.1/CP012721.1 |
| PR06 | SAMN04075681 | PRJNA295544 | GCA_001683195.1 | NZ_CP012723.1/CP012723.1 |
| PR01 | SAMN04075676 | PRJNA295544 | GCA_001683135.1 | NZ_CP012720.1/CP012720.1 |
| PR05 | SAMN04075680 | PRJNA295544 | GCA_001683175.1 | NZ_CP012722.1/CP012722.1 |
| PR07 | SAMN04075682 | PRJNA295544 | GCA_001683215.1 | NZ_CP012724.1/CP012724.1 |
| PR09-1 | SAMN04075684 | PRJNA295544 | GCA_001683255.1 | NZ_CP012726.1/CP012726.1 |
| Parent2 | SAMN04093744 | PRJNA295544 | GCA_001683065.1 | NZ_CP012729.1/CP012729.1 |
| PR08 | SAMN04075683 | PRJNA295544 | GCA_001683235.1 | NZ_CP012725.1/CP012725.1 |
| PR09-4 | SAMN04075685 | PRJNA295544 | GCA_001683275.1 | NZ_CP012727.1/CP012727.1 |
| PR10-4 | SAMN04075686 | PRJNA295544 | GCA_001683295.1 | NZ_CP012728.1/CP012728.1 |

**Table S2. Human respiratory commensal and pathogenic bacterial genome sequences used for the design of *M. amphoriforme* specific baits.** Non-specific baits with BLAST hits to these organisms were removed.

| Accession number | Strain |
|---|---|
| *NC_002929* | *Bordetella pertussis Tohama I* |
| *NZ_CP012981* | *Burkholderia cepacia ATCC 25416, chromosome 1* |
| *NZ_CP012982* | *Burkholderia cepacia ATCC 25416, chromosome 2* |
| *NZ_CP012983* | *Burkholderia cepacia ATCC 25416, chromosome 3* |
| *NZ_LN831026* | *Corynebacterium diphtheriae NCTC11397* |
| NZ_ACEA00000000 | *Eikenella corrodens ATCC 23834* |
| NZ_AJSY00000000 | *Fusobacterium necrophorum* |
| *NC_000907* | *Haemophilus influenzae Rd KW20* |
| *NC_015964* | *Haemophilus parainfluenzae T3T1* |
| *NC_014147* | *Moraxella catarrhalis BBH18* |
| *NC_000962* | *Mycobacterium tuberculosis H37Rv* |
| NC_000908 | *Mycoplasma genitalium G37* |
| NC_013511 | *Mycoplasma hominis ATCC 23114* |
| NZ_ATUH00000000 | *Mycoplasma orale ATCC 23714* |
| NC_000912 | *Mycoplasma pneumoniae M129* |
| NZ_AXZE00000000 | *Mycoplasma salivarium ATCC 23064* |
| NZ_CP007726 | *Neisseria elongata subsp. glycolytica ATCC 29315* |
| NC_014752 | *Neisseria lactamica 020-06* |
| NC_003112 | *Neisseria meningitidis MC58* |
| NC_010729 | *Porphyromonas gingivalis ATCC 33277* |
| NC_014370 | *Prevotella melaninogenica ATCC 25845, chromosome 1* |
| NC_014371 | *Prevotella melaninogenica ATCC 25845, chromosome 2* |
| NZ_AEPE00000000 | *Prevotella oralis ATCC 33269* |
| NZ_ARIR00000000 | *Prevotella veroralis DSM 19559 = JCM 6290* |

| | |
|---|---|
| NC_002516 | *Pseudomonas aeruginosa PAO1* |
| NZ_HG326223 | *Serratia marcescens subsp. marcescens Db11* |
| NC_007795 | *Staphylococcus aureus subsp. aureus NCTC 8325* |
| NC_004461 | *Staphylococcus epidermidis ATCC 12228* |
| NC_013853 | *Streptococcus mitis B6* |
| NC_004350 | *Streptococcus mutans UA159* |
| NC_015291 | *Streptococcus oralis Uo5* |
| NC_003098 | *Streptococcus pneumoniae R6* |
| NC_015875 | *Streptococcus pseudopneumoniae IS7493* |
| NC_002737 | *Streptococcus pyogenes M1 GAS* |
| NZ_CP009913 | *Streptococcus salivarius NCTC 8618* |
| NC_009009 | *Streptococcus sanguinis SK36* |
| NC_010503 | *Ureaplasma parvum serovar 3 str. ATCC 27815* |
| NC_011374 | *Ureaplasma urealyticum serovar 10 str. ATCC 33699* |
| NC_013520 | *Veillonella parvula DSM 2008* |

**Table S3. Summary statistics.** Total reads, mapped reads, fraction of mapped reads, mean depth of coverage (doc), fraction of the genome covered > 15X, for *B. anthracis* and *M. amphoriforme.*

| | *B. anthracis* | *M. amphoriforme* |
|---|---|---|
| median_total_reads | 6413796 | 2989985 |
| max_total_reads | 32708157 | 4247924 |
| min_total_reads | 539623 | 1082454 |
| mean_total_reads | 7624145 | 2918092 |
| median_mapped_reads | 3589129 | 738985 |
| max_mapped_reads | 32291415 | 3056077 |
| min_mapped_reads | 29490 | 10390 |
| mean_mapped_reads | 5442403 | 836116 |
| median_frac_mapped_reads | 0.697 | 0.236 |
| max_frac_mapped_reads | 0.992 | 0.844 |
| min_frac_mapped_reads | 0.023 | 0.003 |
| mean_frac_mapped_reads | 0.611 | 0.294 |
| median_meandoc | 43.19 | 26.95 |
| max_meandoc | 449.39 | 183.05 |
| min_meandoc | 0.17 | 0.08 |
| mean_meandoc | 69.66 | 42.91 |
| iqr_meandoc | 59.96 | 58.93 |
| upper_quartile_meandoc | 74.73 | 64.68 |
| lower_quartile_meandoc | 14.77 | 5.76 |
| median_frac_above15 | 97.3 | 79.2 |
| max_frac_above15 | 100 | 95.8 |
| min_frac_above15 | 0 | 0 |
| mean_frac_above15 | 69.27 | 55.61 |
| iqr_frac_above15 | 55.9 | 88.6 |
| upper_quartile_frac_above15 | 99.8 | 90.8 |
| lower_quartile_frac_above15 | 43.9 | 2.15 |

**Table S4.** Model details for binomial glmm of the effect of *Ct* value, captured library concentration and pooling prior to bait capture on capture efficiency, including an individual-level random effect.

| Results of binomial *glmm* | Estimate | Std. Error | z value | Pr(>\|z\|) |
|---|---|---|---|---|
| (Intercept) | 9.66832 | 0.78245 | 12.357 | < 2e-16 *** |
| max_ct | -0.35679 | 0.02788 | -12.799 | < 2e-16 *** |
| cap_lib_conc | 0.05805 | 0.01231 | 4.715 | 2.42e-06 *** |
| pooled_yesorno | -1.31290 | 0.27892 | -4.707 | 2.42e-06 *** |

| *drop1* output: | npar | AIC | LRT | Pr(Chi) |
|---|---|---|---|---|
| <none> | | 4451.8 | | |
| max_ct | 1 | 4541.5 | 110.716 | < 2.2e-16 *** |
| cap_lib_conc | 1 | 4451.5 | 20.726 | 5.299e-06 *** |
| pooled_yesorno | 1 | 4451.4 | 20.666 | 5.468e-06 *** |

| *MuMIn* output: | | R2m | R2c |
|---|---|---|---|
| | theoretical | 0.6124126 | 0.9999999 |
| | delta | 0.6124126 | 0.9999999 |

9. **Hartig F**. *DHARMa: residual diagnostics for hierarchical (multi-level/mixed) regression models*. https://cran.r-project.org/web/packages/DHARMa/vignettes/DHARMa.html.

10. **Wilke CO**. *cowplot – Streamlined plot theme and plot annotations for ggplot2*. https://wilkelab.org/cowplot/.
